# Bias in miRNA enrichment analysis related to gene functional annotations

**DOI:** 10.1101/2021.08.16.456527

**Authors:** Konstantinos Zagganas, Thanasis Vergoulis, Georgios K. Georgakilas, Spiros Skiadopoulos, Theodore Dalamagas

## Abstract

**Background:** miRNA functional enrichment is a type of analysis that is used to predict which biological functions may be affected by a group of miRNAs or validate whether a list of dysregulated miRNAs are linked to a diseased state. The standard method for functional enrichment analysis uses the hypergeometric distribution to produce p-values, depicting the strength of the association between a group of miRNAs and a biological function. However, in 2015, it was shown that this approach suffers from a bias related to miRNA targets produced by target prediction algorithms and a new randomization test was proposed to alleviate this issue.

**Results:** We demonstrate the existence of another previously unreported underlying bias which affects gene annotation data sets; additionally, we show that the statistical measure used for the established randomization test is not sensitive enough to account for it. In this context, we show that the use of Jaccard coefficient (an alternative statistical measure) is able to alleviate the aforementioned issue.

**Conclusions:** In this paper, we illustrate the existence of a new bias affecting the miRNA functional enrichment analysis. This bias makes Fisher’s exact test unsuitable for miRNA functional enrichment analyses and there is also a need to adjust the established unbiased test accordingly. We propose the use of a modified version of the established test and in order to facilitate its use, we introduce a novel unbiased miRNA enrichment analysis tool that implements the proposed method. At the same time, by leveraging bit vectors, our tool guarantees fast and scalable execution.

**Availability:** All datasets used in the experiments throughout this paper are openly accessible on Zenodo (https://doi.org/10.5281/zenodo.5175819).

## 1 Background

miRNAs are short, non-coding RNA molecules (~23 nucleotides long) which are important posttranscription gene expression regulators. They exert their function by binding to mRNAs or long non-coding RNAs (lncRNAs). This “silences” RNAs in two ways: (1) by facilitating the degradation of RNAs and (2) by inhibiting ribosomes from synthesising proteins. Since gene expression leads to protein production and thus, is an integral part of cellular structure and function, this means that, by regulating gene expression, miRNAs can potentially affect a multitude of biological pathways. Due to their significant importance, computational analyses that attempt to investigate miRNA functions and their involvement in various biological pathways have been popular, in recent years. One such type of analysis in the so-called *miRNA functional enrichment analysis* (e.g., [22]). Generally, given a collection of miRNAs and a biological function, miRNA functional enrichment analysis usually involves the following steps: (1) retrieve a list of all genes targeted by the group of miRNAs, which can either be the union or the intersection of the genes targeted by each individual miRNA in the group, (2) retrieve the list of genes that participate in the biological function and (3) perform a statistical test, usually Fisher’s exact test [6], to calculate a p-value that indicates the strength of the association between the miRNA group and the biological function (i.e. how likely is the biological function to be affected by the group).

However, in 2015, it was demonstrated by Bleazard *et al*. [3], that the standard method of miRNA functional enrichment analysis is not suitable for such analyses, since it provides highly unspecific results. Briefly, Bleazard *et al*. illustrated that miRNA functional enrichment analysis based on target prediction algorithms is strongly biased and leads to the identification of unrelated biological processes. The mechanics responsible for this bias are not yet fully understood due to the fact that miRNA target prediction is far from being a solved problem in Bioinformatics ([16, 1]). Yet, some contributing factors can be identified.

More specifically, one of them is the limited amount of validated positive miRNA:target interactions, and more importantly the virtually non-existent validated negative interactions, which severely affect the training of robust and highly accurate target prediction algorithms [13]. To make matters worse, most target prediction algorithms have been trained on seed-enriched data sets with features extracted from the sequence surrounding the seed, even though recent evidence shows that non-seed-based interactions are common in miRNA-mediated gene expression regulation [14, 10]. The inefficient selection of negative samples for the training process is often unavoidable due to the small number of validated negative miRNA:target interactions in the literature.

To make matters more complicated, the process of experimentally validating miRNA binding sites is frequently driven by target prediction algorithms. Life scientists often apply a target prediction algorithm on a gene under study, select the top target and carry out the validation process. Negative results are usually not reported while the published positive interactions are inevitably enriched in seed-based binding types. The whole back and forth between miRNA target prediction results and experimentally-driven interaction validation results in a vicious circle that inflates miRNA functional enrichment analyses with false positive target genes. This, inevitably makes the bias strong enough to invalidate the assumption made by the hypergeometric distribution, that genes are targeted by miRNAs in a uniform fashion. Moreover, Bleazard *et. al* illustrated that statistically significant results with the standard method did not remain so, after correcting for bias.

In order to eliminate the bias affecting the standard overrepresentation analysis, Bleazard *et al*. proposed a new method to perform miRNA functional enrichment analysis, that involves a randomization test. This test moves the analysis to the miRNA level instead of the gene level and it consists the following steps:

1. Given a miRNA group of interest calculate a statistical measure relevant to the problem.
2. Create 1 million randomly assembled miRNA groups with the same size as the group of interest and for each of them calculate the same statistical measure.
3. The empirical p-value is then defined as the proportion of randomly assembled miRNA groups that present a better statistical behaviour compared to the behaviour of the group of interest.

The statistical measure used is called *GO term overlap* [3] and it is defined as the proportion of genes targeted by a group of miRNAs, that are also members of a specific GO category. Let *A* be the set of genes targeted by the group and *B* be the set of genes that participate in the GO category. Then the GO term overlap, to which we are going to refer as *left-sided-overlap* from here on, is more formally defined as:

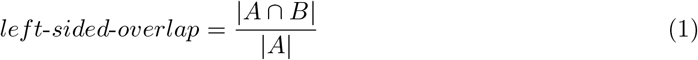

Intuitively we expect that the left-sided overlap accounts for the bias introduced by target prediction algorithms, which is the main focus of [3]. This is done by dividing the size of the overlap between the set of targets and the GO category by the number of the targets.

In this paper we demonstrate that when target-prediction bias is removed, by using experimentally validated miRNA targets, then a new bias, related to gene annotations, is made visible. Additionally, we illustrate that this occurs due to a bias affecting the gene-to-biological-function annotations and is not accounted for by the unbiased enrichment analysis proposed by Bleazard *et al*., resulting in reduced sensitivity to false negatives. To alleviate this issue, we introduce a new statistical measure that increases the sensitivity of the unbiased enrichment analysis. This measure accounts for both sources of bias (predicted interactions and gene annotations) and reduces false negative results when either predicted or experimentally validated targets are used. Finally, leveraging the lack of tools performing the proposed, modified, unbiased miRNA functional enrichment analysis, we introduce BUFET2, that calculates p-values using both statistical measures. BUFET2 is available as source code, Docker image, and a REST API service.

## 2 Methods

### 2.1 Investigating experimentally validated miRNA targets

The aim of this study, is to show that a bias affecting miRNA functional enrichment analysis still persists even when experimentally validated miRNA targets are used, instead of predicted targets. To do this, we follow an approach similar to Bleazard *et al*. to show that the empirical distribution of the overlaps does not match with the expected hypergeometric distribution. First, we selected miRNAs dysregulated in multiple sclerosis, from Table 5 in [15] (see supporting information) and gene ontology (GO) annotations from Ensembl [23] version 100. Also we retrieved disease-to-gene associations from DisGeNET v7 [18], as well as pathway data from KEGG [11]. Moreover, we used the latest version of miRTarBase [4] at the time this article is written, in order to get a list of experimentally validated miRNA targets. We selected 14 of the dysregulated miRNAs that exist in the data set.

Initially, we randomized at the miRNA level, by creating 1 million randomly assembled miRNA groups containing 14 miRNAs each. For each of these groups, we calculated the gene members of each GO category in the data set, that are also targeted by the miRNAs in the group and created a histogram. On the same graph, we plotted the expected hypergeometric distribution for the overlaps, given the number of targeted/non-targeted genes and the number of genes belonging/not-belonging to the same GO term. The results of this experiment are presented in Section “Comparison of Empirical and Hypergeometric distribution”.

### 2.2 Introducing the use of the Jaccard coefficient

Based on the definition of the left-sided overlap, we can see that it is the proportion of targets that also belong to the gene class. Intuitively, this suggests that the left-sided overlap is designed to remove the bias stemming from target prediction, because the intersection is normalized using the total number of genes being targeted by the miRNA group. To eliminate the bias related to the size of each gene class, we are proposing the use of the Jaccard coefficient [8] as a metric for the randomization test:

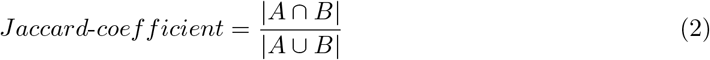

We argue that normalization taking into account the total number of genes involved between a gene class and a miRNA group will increase the overall sensitivity of the test and it should be the preferred method regarding miRNA functional enrichment analysis.

With this in mind, in Section “Left-sided overlap vs Jaccard coefficient, we retrieved published collections of dysregulated miRNAs in diseases like Alzheimer’s and several cancer types, in order to show that accounting only for the bias from miRNA target prediction leads to decreased sensitivity and consequently to false negative results. To prove this, we compared the p-values produced from using the left-sided-overlap and the Jaccard coefficient for specific pathways related to Alzheimer’s disease and cancer. Moreover, in Section “Randomization test metric vs type of type of miRNA targets”, we performed the same experiment for the lists used in the “Left-sided overlap vs Jaccard coefficient” section, using miRNA targets from microT-CDS [19] and compared the p-values produced using the left-sided overlap as well as Jaccard coefficient.

## 3 Results

### 3.1 Comparison of Empirical and Hypergeometric distribution

In this section we demonstrate that even when the bias from miRNA target prediction is removed, the hypergeometric test still presents decreased robustness, due to a bias related to gene class data. For this reason, we used gene annotation data from GO to illustrate that the expected hypergeometric distribution differs from the empirical distributions of the overlaps between a gene class and a miRNA group.

In order to compare the two distributions, we first calculated the number of targets for the 14 miRNAs in the group. We found that 3106 out of a total of 15064 genes are indicated as targets in the set of interactions. Then, given the size of each GO category and the genes not targeted by our original miRNA group, we calculated the expected hypergeometric distribution following a similar approach to the one followed by Bleazard *et al*. The next step was to estimate the empirical distribution. First, we created 1 million randomly assembled miRNA groups of size 14 and for each of those miRNA groups we calculated the size of the intersection between the targets of the group and each GO category. Finally, to facilitate comparison of the two distributions, two separate histograms were created and plotted on the same graph.

Considering that our data set for GO consists of 18686 categories, we selected indicative examples, for the diverse behaviour of the two distributions. These examples include, among others, the three domains of GO (biological process, molecular function, cellular component) and ion transport (since it was also used by Bleazard *et al*.).The results are illustrated in Figure 1.

**Figure 1:**
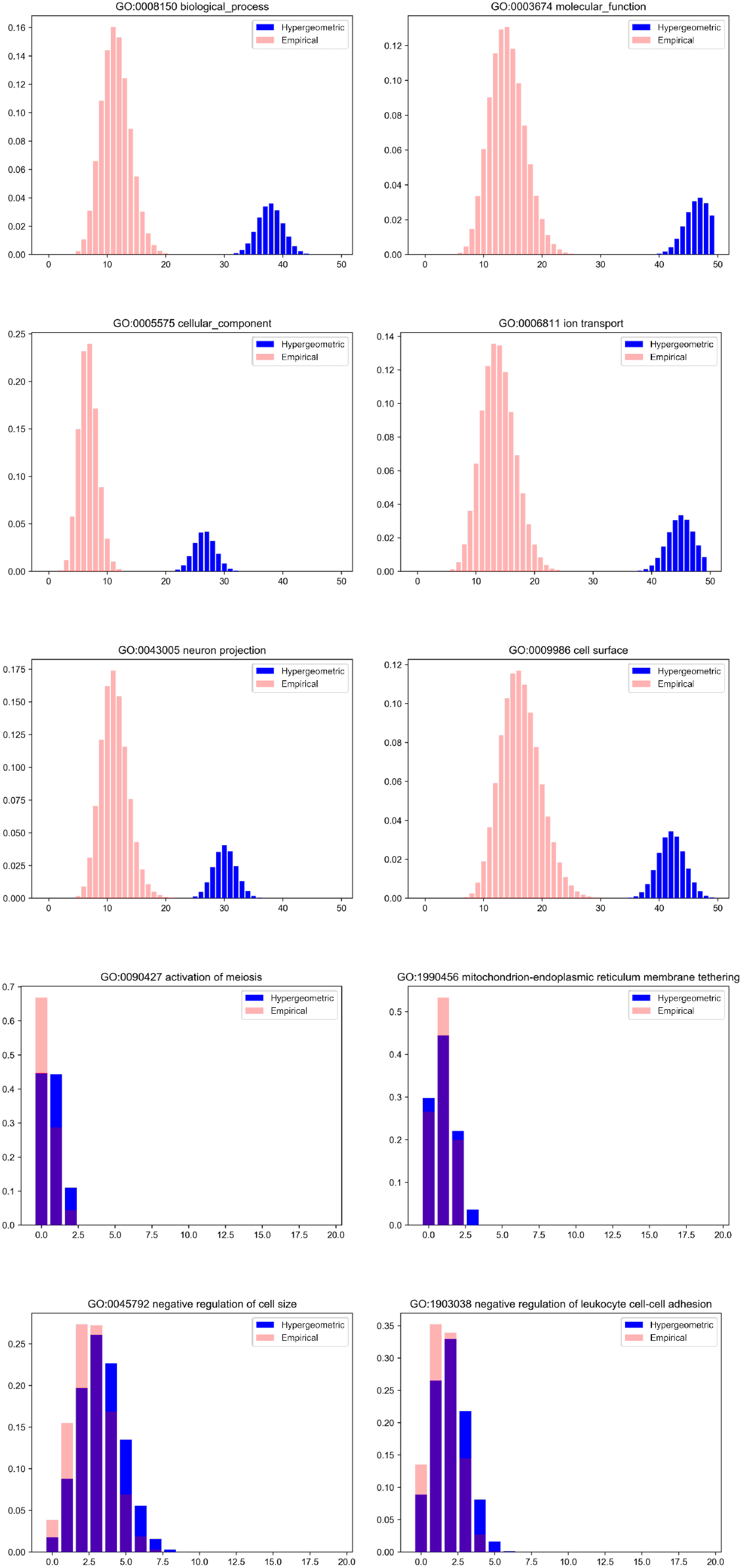
Hypergeometric vs empirical distributions for different GO categories

Immediately, it becomes evident that there are categories for which the two distributions match and categories for which there is a clear mismatch between them. Categories, which are higher in the GO hierarchy seem to have a larger divergence between the distributions. On the other hand, categories, that describe more specific biological functions, tend to significantly overlap with the hypergeometric distribution. Moreover, the categories that present the larger mismatch seem to be those, that contain a large number of genes. This provides the intuition, that maybe, the size of the GO category is related to the mismatch between the distributions. In Table 1 we demonstrate the sizes of each of the GO categories shown in Figure 1.

**Table 1:** Number of genes in each of the GO categories in Figure 1.

| Accession number | Name | Size |
| --- | --- | --- |
| GO:0008150 | biological_process | 570 |
| GO:0003674 | molecular_function | 707 |
| GO:0005575 | cellular_component | 401 |
| GO:0006811 | ion transport | 679 |
| GO:0043005 | neuron projection | 453 |
| GO:0009986 | cell surface | 634 |
| GO:0090427 | activation of meiosis | 2 |
| GO:1990456 | mitoch. endopl. reticulum memb. tethering | 3 |
| GO:0045792 | neg. regulation of cell size | 10 |
| GO:1903038 | neg. reg. of leukocyte cell-cell adh. | 6 |

Given the results in Table 1, it seems that the mismatch is indeed more prominent as the size of the category increases. In order to quantify and investigate this effect, we designed the following experiment. First, we define as *distance* between the distributions the horizontal distance (number of genes in common with the GO category) between the maximum values of each distribution. Then, using this definition, for all GO categories in the data set, we evaluate the distance between the two distributions in relation to the category size. The result of this experiment can be seen in Figure 2.

**Figure 2:**
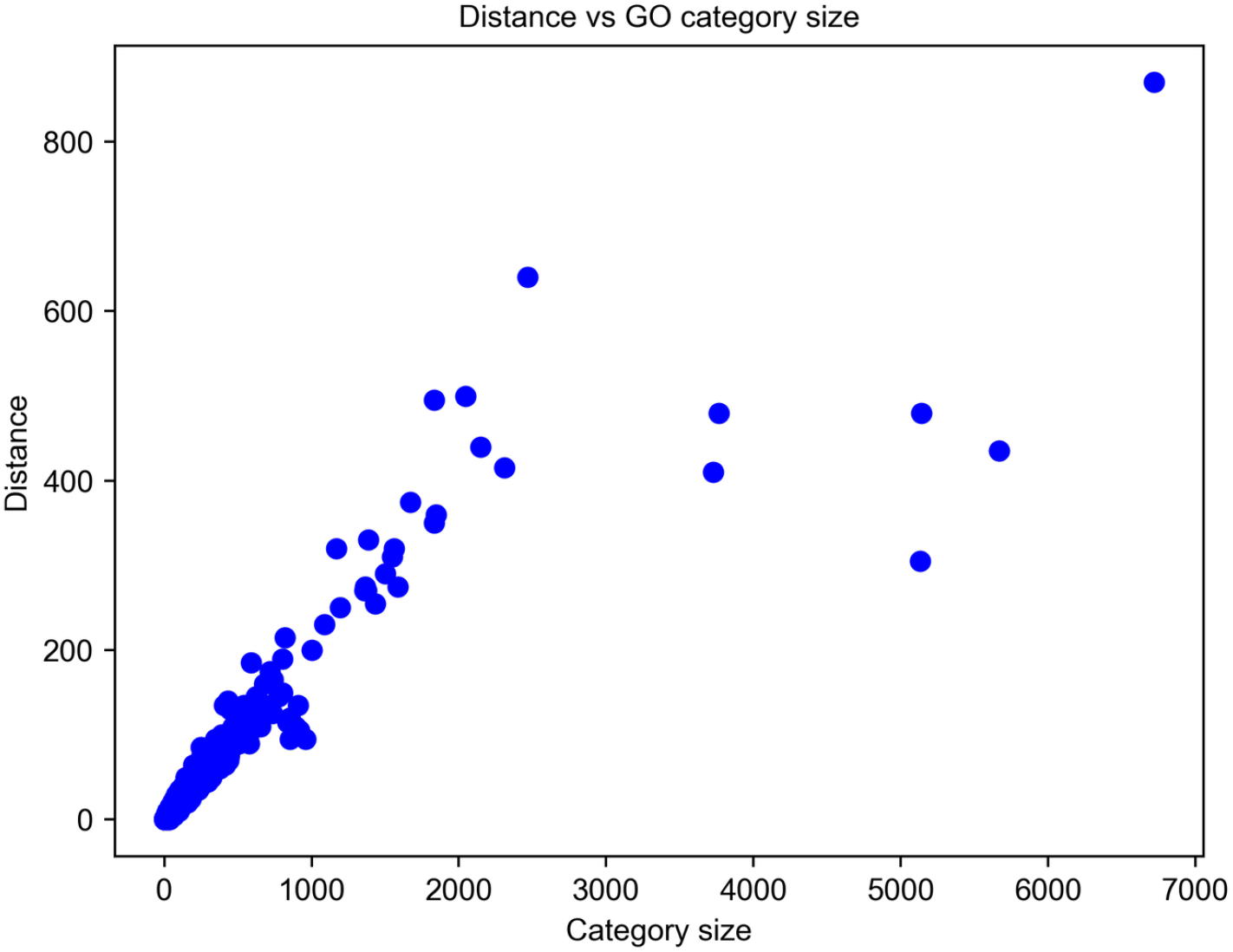
Distance between hypergeometric and empirical distribution vs GO category size

We can clearly observe that, barring a few outliers, in general, the greater the size of a GO category, the larger the distance is and this translates to a larger mismatch between the two distributions. Given the hierarchical structure of the Gene Ontology, it is easy to understand why this phenomenon exists: the genes in a GO category are also contained in a category which is higher in the hierarchy (e.g. neuron projection, cell projection and cellular component) and this means that the genes in a category higher in the tree are not picked uniformly and thus they do not satisfy the assumptions of the hypergeometric distribution. This, essentially introduces a bias that is more prominent, the higher up from the branches of the tree one goes. To check whether this effect can also be observed with DisGeNET and KEGG, we performed the same experiment using these data sets. The results can be seen in Figure 3 and Figure 4 respectively. Evidently, the aforementioned effect is even more pronounced for DisGeNET and the relationship between the disease size and the distance between the two distributions. This can maybe be explained by the fact that the text mining tools used to compile the database, utilize structured vocabularies and ontologies [18]. Thus, the hierarchy existing between the diseases introduces the same bias as seen for GO. Regarding KEGG, the same effect is also observed. This could maybe be attributed to complex interactions between genes in pathways that are not specified in the data set. An example of this could be gene co-expression or other more complex effects that lead to genes being included in a pathway.

**Figure 3:**
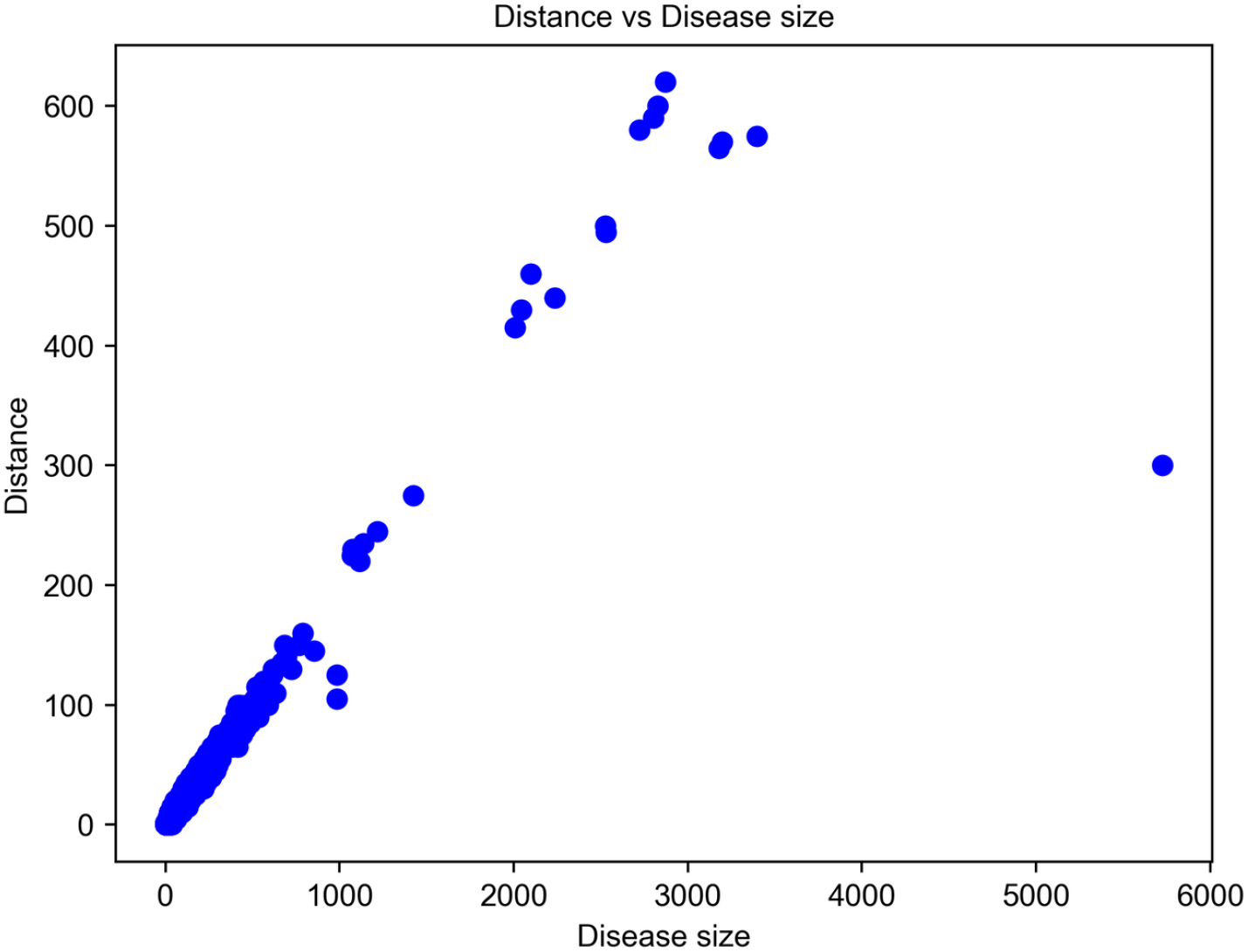
Distance between hypergeometric and empirical distribution vs disease size

**Figure 4:**
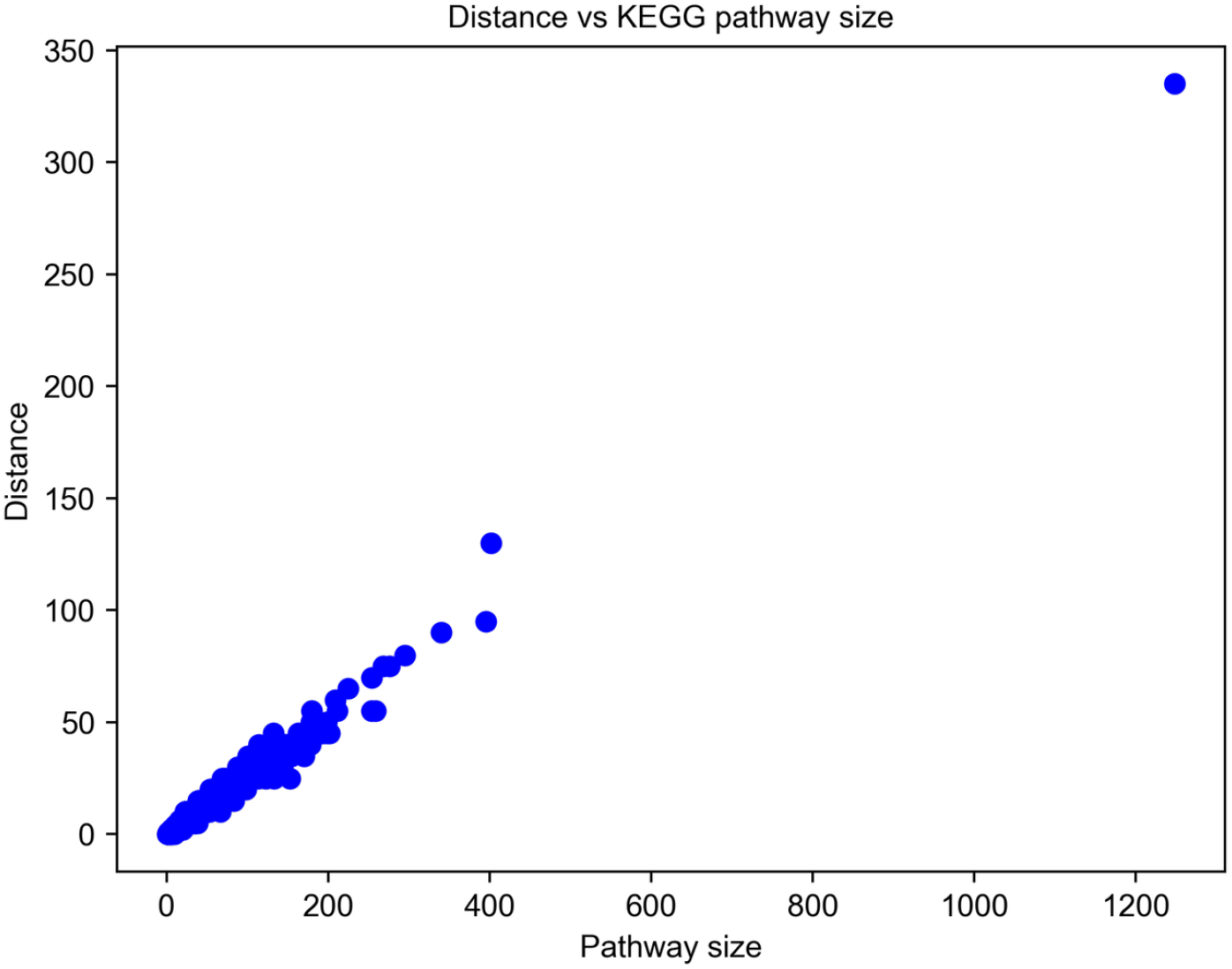
Distance between hypergeometric and empirical distribution vs size of KEGG pathway

In conclusion, we showed that for all three data sets there is a bias, which is clearly a product of the gene annotation structure, affecting the miRNA functional enrichment analysis.

### 3.2 Left-sided overlap vs Jaccard coefficient

In the previous section, we demonstrated that there is a clear bias related to the structure of the gene annotations. In this section we show that the left-sided overlap does not account for this bias, highlighting the need to use another, more sensitive metric, namely the Jaccard coefficient. For this reason, we compared the performance of the two metrics using four, published lists of miRNAs: one for Alzheimer’s disease [2] one for non-small-cell lung cancer [26], one for breast cancer [20] and one for gastric cancer [9]. These lists are provided as supporting information.

Furthermore, we utilized two modules of the KEGG database, namely PATHWAY and DISEASE, since DISEASE provides pathways associated with each of the four diseases. It stands to reason that if we provide the list of miRNAs for each disease to the randomization test as input, at least some (if not all) of the pathways related to each disease should be marked as significant. Apart from the pathways related specifically to each disease, we will present other pathways commonly associated with cancers (for the three cancer lists). Each experiment was performed 10 times and in Tables 2, 3, 4 and 5 we present the average of each p-value along with the standard deviation for the 10 experiments.

**Table 2:**
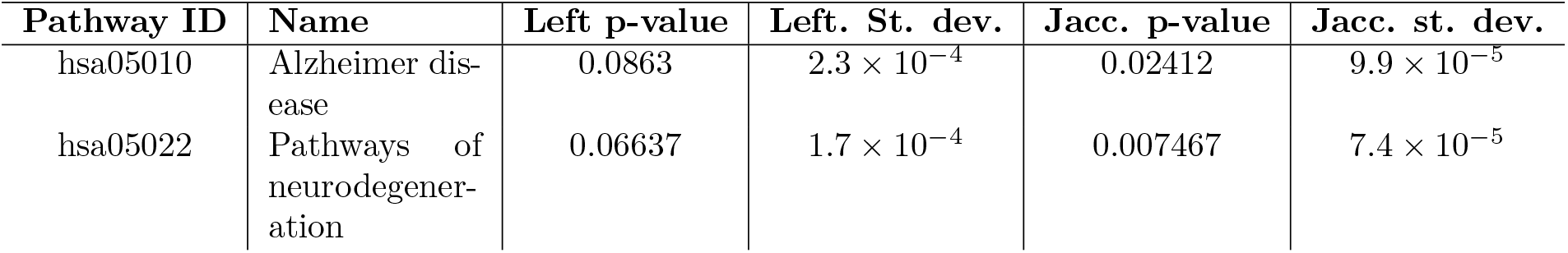
Left-sided and Jaccard p-values for Alzheimer’s disease using miRTarBase.

| Pathway ID | Name | Left p-value | Left. St. dev. | Jacc. p-value | Jacc. st. dev. |
| --- | --- | --- | --- | --- | --- |
| hsa05010 | Alzheimer dis-<br>ease | 0.0863 | $2.3 \times 10^{-4}$ | 0.02412 | $9.9 \times 10^{-5}$ |
| hsa05022 | Pathways of<br>neurodegener-<br>ation | 0.06637 | $1.7 \times 10^{-4}$ | 0.007467 | $7.4 \times 10^{-5}$ |

**Table 3:** Left-sided and Jaccard p-values for Non-small-cell lung cancer using miRTarBase.

| Pathway ID | Name | Left p-value | Left St. dev. | Jacc. p-value | Jacc. st. dev. |
| --- | --- | --- | --- | --- | --- |
| hsa05223 | Non-small-cell<br>lung cancer | 0.2606 | $3.9 \times 10^{-4}$ | 0.229 | $3.5 \times 10^{-4}$ |
| hsa05200 | Pathways in<br>cancer | 0.01745 | $2.1 \times 10^{-4}$ | 0.0005191 | $1.9 \times 10^{-5}$ |
| hsa05206 | MicroRNAs in<br>cancer | 0.0867 | $2.6 \times 10^{-4}$ | 0.03432 | $1.5 \times 10^{-4}$ |
| hsa05205 | Proteoglycans<br>in cancer | 0.06989 | $1.5 \times 10^{-4}$ | 0.03196 | $1.5 \times 10^{-4}$ |

**Table 4:** Left-sided and Jaccard p-values for breast cancer using miRTarBase.

| Pathway ID | Name | Left p-value | Left St. dev. | Jacc. p-value | Jacc. st. dev. |
| --- | --- | --- | --- | --- | --- |
| hsa05224 | Breast cancer | 0.00436 | $8.6 \times 10^{-5}$ | 0.002436 | $5 \times 10^{-5}$ |
| hsa05200 | Pathways in<br>cancer | 0.000223 | $1.7 \times 10^{-5}$ | 0.0000164 | $3.7 \times 10^{-6}$ |
| hsa05206 | MicroRNAs in<br>cancer | 0.0001501 | $1.6 \times 10^{-5}$ | 0.0000107 | $3.3 \times 10^{-6}$ |
| hsa04020 | Calcium<br>signaling<br>pathway | 0.07019 | $1.7 \times 10^{-4}$ | 0.04543 | $2.1 \times 10^{-4}$ |

**Table 5:** Left-sided and Jaccard p-values for gastric cancer using miRTarBase.

| Pathway ID | Name | Left p-value | Left St. dev. | Jacc. p-value | Jacc. st. dev. |
| --- | --- | --- | --- | --- | --- |
| hsa05226 | Gastric cancer | 0.03668 | $2 \times 10^{-4}$ | 0.02293 | $1.3 \times 10^{-4}$ |
| hsa05200 | Pathways in cancer | 0.01216 | $1.5 \times 10^{-4}$ | 0.0005839 | $1.7 \times 10^{-5}$ |
| hsa05206 | MicroRNAs in cancer | 0.083 | $3.8 \times 10^{-4}$ | 0.03825 | $2.4 \times 10^{-4}$ |

Tables 2, 3, 4 and 5 immediately suggest that using the Jaccard coefficient as a metric increases the sensitivity of the test to false negatives and leads to more accurate results. This is particularly evident with Alzheimer’s disease, where the “Alzheimer’s disease” pathway is not marked as statistically significant by the left-sided p-value. Moreover, even when both methods suggest that a pathway is significant, the Jaccard p-value is smaller, denoting a stronger association of the miRNA group with the pathway. The complete list of results is provided as additional files.

### 3.3 Randomization test metric vs type of type of miRNA targets

In this section we perform the same experiment as in the previous section, but this time, instead of experimentally validated results, we utilize miRNA targets predicted using microT-CDS, retrieved from the MR-microT[12] online application using a prediction score threshold of 0.8.

The results (see Additional Files) paint a picture similar to the previous section. As an example we present in Table 6 some indicative results for non-small-cell lung cancer. It becomes evident that the use of predicted targets leads to more false negative results than those procured from the utilization of validated targets for both metrics. However, even in this case the Jaccard p-value seems to present a larger sensitivity.

**Table 6:**
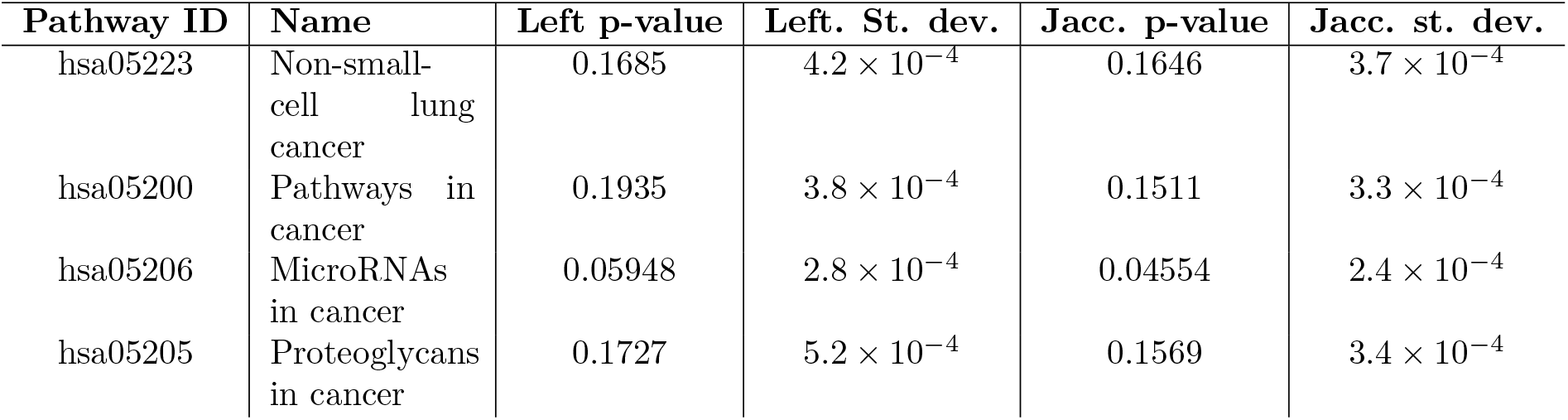
Left-sided and Jaccard p-values for Non-small-cell lung cancer using microT.

## 4 Discussion

Issues regarding overrepresentation analysis and the hierarchy of Gene Ontology annotations have been mentioned as far back as 2003 [5, 25]. Bleazard *et al*. also point out that there may be underlying correlations between targeting of processes and the hierarchical GO structure. In the “Comparison of Empirical and Hypergeometric distribution” section we demonstrated that there is a definite bias related to the hierarchical structure of the GO and that this bias also affects other gene annotation data sets like KEGG PATHWAY and DisGeNET. This suggests that Fisher’s exact test does not present enough robustness for miRNA functional enrichment analyses, even when the bias from miRNA prediction algorithms is eliminated through the utilization of experimentally validated miRNA targets. Despite its drawbacks, however, the standard method is still very popular in published studies [7].

Furthermore, the “Left-sided overlap vs Jaccard coefficient” and “Randomization test metric vs type of type of miRNA targets” sections clearly demonstrate that the use of the Jaccard coefficient leads to a more sensitive test that produces less false negative results. This denotes that the same bias affects the analysis regardless of the type of interactions used. On the other hand, it becomes evident that the enrichment analysis using miRNA predicted targets is not as sensitive as the one using experimentally validated targets. This is expected and can be attributed to the fact that prediction algorithms produce hundreds or thousands of targets for each miRNA and the results contain a large number of false positive interactions [17]. This implies that genes in categories or diseases at the bottom of the hierarchy (more specific) are being targeted by a lot of miRNAs, even if these are not true interactions. This translates to many randomly assembled groups that often present a better overlap than the sample of interest and consequently, the sample of interest loses its status of being among the 5% of miRNA groups in the empirical distribution (i.e. p-value≥0.05)

We should also note here an important observation regarding empirical p-values and randomization tests. More specifically, in our case, the standard deviation between experiments is at least two orders of magnitude smaller than the p-value significance threshold (0.05), for both types of overlaps. This denotes that p-values produced by the 10 repetitions of the same experiment do not present with a large variation in significance. In other words, if a gene class is marked as significant by one of the 10 repetitions, then the magnitude of the standard deviation implies that the statistical significance of the gene class will be preserved across every repetition of the same experiment. Consequently, no matter how many times one runs the experiment with the same input, the produced p-values are not expected to be qualitatively altered. In general, this shows that 1 million random groups are sufficient for this randomization test. It should be mentioned here that the aim of randomization tests is to model the empirical distribution of the data in order to reach conclusions. As a result, the accuracy of the empirical p-values produced depends mainly on the number of random samples used for the randomization test. A small number of random samples is not enough to model the empirical p-values and the results can present a high variation between two repetitions of the same experiment. On the other hand, a very large number of random samples is not necessary since it can consume a large number of compute resources and lead to significantly large execution times; this is because it does not really contribute much to the results, considering that a smaller number of samples can accurately predict whether the sample of interest presents a p-value smaller than 0.05. Thus, such randomization tests require a balance between too few and too many random samples and sufficient accuracy can be reached when the results between two repetitions of the same experiment do not vary in a way that changes whether a result is significant or not.

Motivated by the decreased sensitivity of tools that estimate the empirical distribution utilizing the left-sided overlap (like empiricalGO [3] and BUFET [24]), we introduce BUFET2. BUFET2 is designed to provide two p-values for the same randomization test that are produced using the leftsided overlap and the Jaccard coefficient as a statistical measure, respectively. BUFET2, inspired by BUFET, harnesses the power of special data structures, called bit vectors to provide scalable performance and produce results in a matter of minutes for the specified accuracy of 1 million random miRNA groups. To our knowledge, no other tool exists that implements this type of unbiased miRNA enrichment analysis.

At the same time, BUFET2 comes with a new, novel *reverse search module* that allows researchers to discover statistically significant microRNAs associated with a gene class (GO term, disease, pathway, etc), in almost real time. The module takes a gene class as input and randomizes at the miRNA level; however, the randomization is not a Monte Carlo approach, but rather an exhaustive one, since the number of known miRNAs is relatively small (~ 2500). It should be noted here that the reverse search module also estimates both p-values. Moreover, the advantage of our software compared to other reverse search approaches, like miRPath v.3 [21], is that it allows users to provide custom interaction and gene annotation data sets, making our approach significantly more flexible.

Finally, both modules are also publicly available via a REST API^1^ which facilitates programmatic access to the analysis. The API is implemented using Python^2^ and it is documented using OpenAPI and Swagger^3^. Finally the source code^4^ (openly accessible under a GPL v3.0 licence) has been packaged in a Docker image^5^ to facilitate easier execution of the code without the need for compilation.

## 5 Conclusion

Concluding, in this paper we demonstrated that when the bias from miRNA target prediction algorithms is removed, then another bias, affecting gene classes appears. This bias has a clear relation to the size of each class and it might be related to a hierarchical structure or other nonspecific reasons. Additionally, we suggest the use of the Jaccard coefficient, which seems to be more appropriate as a randomization test metric in all cases (predicted and experimentally validated targets). Furthermore, all data sets used in the Results section have been uploaded to Zenodo (see Abstract) to facilitate experiment reproduction. Finally, we introduced BUFET2 a novel software that calculates p-values using the empirical distribution and at the same time utilizing both the leftsided overlap as well as the Jaccard coefficient as metrics to increase the accuracy of the analysis.

## Footnotes

1 https://bufet2.imsi.athenarc.gr/

2 https://www.python.org/

3 https://swagger.io/specification/

4 https://github.com/athenarc/bufet2

5 https://hub.docker.com/r/diwis/bufet2

## Notes

https://zenodo.org/record/5175819#.YRq4TO1RW6s

### Competing Interest Statement

The authors have declared no competing interest.

## References

[1] Panagiotis Alexiou, Manolis Maragkakis, Giorgos L. Papadopoulos, Martin Reczko, and Artemis G. Hatzigeorgiou. Lost in translation: an assessment and perspective for computational microRNA target identification. Bioinformatics, 25(23):3049–3055, 09 2009.

[2] Francesco Angelucci, Katerina Cechova, Martin Valis, Kamil Kuca, Bing Zhang, and Jakub Hort. Micrornas in alzheimer’s disease: Diagnostic markers or therapeutic agents? Frontiers in Pharmacology, 10:665, 2019.

[3] Thomas Bleazard, Janine A Lamb, and Sam Griffiths-Jones. Bias in microRNA functional enrichment analysis. Bioinformatics, 31(10):1592–1598, 01 2015.

[4] Chih-Hung Chou, Sirjana Shrestha, Chi-Dung Yang, Nai-Wen Chang, Yu-Ling Lin, Kuang-Wen Liao, Wei-Chi Huang, Ting-Hsuan Sun, Siang-Jyun Tu, Wei-Hsiang Lee, Men-Yee Chiew, Chun-San Tai, Ting-Yen Wei, Tzi-Ren Tsai, Hsin-Tzu Huang, Chung-Yu Wang, Hsin-Yi Wu, Shu-Yi Ho, Pin-Rong Chen, Cheng-Hsun Chuang, Pei-Jung Hsieh, Yi-Shin Wu, Wen-Liang Chen, Meng-Ju Li, Yu-Chun Wu, Xin-Yi Huang, Fung Ling Ng, Waradee Buddhakosai, Pei-Chun Huang, Kuan-Chun Lan, Chia-Yen Huang, Shun-Long Weng, Yeong-Nan Cheng, Chao Liang, Wen-Lian Hsu, and Hsien-Da Huang. miRTarBase update 2018: a resource for experimentally validated microRNA-target interactions. Nucleic Acids Research, 46(D1):D296–D302, 11 2017.

[5] Sorin Drăghici, Purvesh Khatri, Rui P. Martins, G.Charles Ostermeier, and Stephen A. Krawetz. Global functional profiling of gene expression. Genomics, 81(2):98 – 104, 2003.

[6] R. A. Fisher. Statistical Methods for Research Workers, pages 66–70. Springer New York, New York, NY, 1992.

[7] Adrian Garcia-Moreno and Pedro Carmona-Saez. Computational methods and software tools for functional analysis of mirna data. Biomolecules, 10(9):1252, Aug 2020.

[8] John M. Hancock. Jaccard Distance (Jaccard Index, Jaccard Similarity Coefficient). American Cancer Society, 2014.

[9] Ning-Bo Hao, Ya-Fei He, Xiao-Qin Li, Kai Wang, and Rui-Ling Wang. The role of mirna and lncrna in gastric cancer. Oncotarget, 8(46):81572–81582, 2017.

[10] Aleksandra Helwak, Grzegorz Kudla, Tatiana Dudnakova, and David Tollervey. Mapping the human mirna interactome by clash reveals frequent noncanonical binding. Cell, 153(3):654–665, 2013.

[11] Minoru Kanehisa and Susumu Goto. KEGG: Kyoto Encyclopedia of Genes and Genomes. Nucleic Acids Research, 28(1):27–30, 01 2000.

[12] Ilias Kanellos, Thanasis Vergoulis, Dimitris Sacharidis, Theodore Dalamagas, Artemis Hatzigeorgiou, Stelios Sartzetakis, and Timos Sellis. Mr-microt: A mapreduce-based microrna target prediction method. In Proceedings of the 26th International Conference on Scientific and Statistical Database Management, SSDBM ’14, New York, NY, USA, 2014. Association for Computing Machinery.

[13] Dimitra Karagkouni, Maria D Paraskevopoulou, Serafeim Chatzopoulos, Ioannis S Vlachos, Spyros Tastsoglou, Ilias Kanellos, Dimitris Papadimitriou, Ioannis Kavakiotis, Sofia Maniou, Giorgos Skoufos, Thanasis Vergoulis, Theodore Dalamagas, and Artemis G Hatzigeorgiou. DIANA-TarBase v8: a decade-long collection of experimentally supported miRNA–gene interactions. Nucleic Acids Research, 46(D1):D239–D245, 11 2017.

[14] Gabriel B. Loeb, Aly A. Khan, David Canner, Joseph B. Hiatt, Jay Shendure, Robert B. Darnell, Christina S. Leslie, and Alexander Y. Rudensky. Transcriptome-wide mir-155 binding map reveals widespread noncanonical microrna targeting. Molecular Cell, 48(5):760–770, Dec 2012.

[15] Eiman MA Mohammed. Environmental influencers, microrna, and multiple sclerosis. Journal of Central Nervous System Disease, 12:1179573519894955, 2020.

[16] Sarah Peterson, Jeffrey Thompson, Melanie Ufkin, Pradeep Sathyanarayana, Lucy Liaw, and Clare Bates Congdon. Common features of microrna target prediction tools. Frontiers in Genetics, 5:23, 2014.

[17] Natalia Pinzón, Blaise Li, Laura Martinez, Anna Sergeeva, Jessy Presumey, Florence Apparailly, and Hervé Seitz. microrna target prediction programs predict many false positives. Genome Research, 27(2):234–245, 2017.

[18] Janet Piñero, Juan Manuel Ramírez-Anguita, Josep Saüch-Pitarch, Francesco Ronzano, Emilio Centeno, Ferran Sanz, and Laura I Furlong. The DisGeNET knowledge platform for disease genomics: 2019 update. Nucleic Acids Research, 48(D1):D845–D855, 11 2019.

[19] Martin Reczko, Manolis Maragkakis, Panagiotis Alexiou, Ivo Grosse, and Artemis G. Hatzigeorgiou. Functional microRNA targets in protein coding sequences. Bioinformatics, 28(6):771–776, 01 2012.

[20] Eleni van Schooneveld, Hans Wildiers, Ignace Vergote, Peter B. Vermeulen, Luc Y. Dirix, and Steven J. Van Laere. Dysregulation of micrornas in breast cancer and their potential role as prognostic and predictive biomarkers in patient management. Breast Cancer Research, 17(1):21, Feb 2015.

[21] Ioannis S. Vlachos, Konstantinos Zagganas, Maria D. Paraskevopoulou, Georgios Georgakilas, Dimitra Karagkouni, Thanasis Vergoulis, Theodore Dalamagas, and Artemis G. Hatzigeorgiou. DIANA-miRPath v3.0: deciphering microRNA function with experimental support. Nucleic Acids Research, 43(W1):W460–W466, 05 2015.

[22] Carrie Wright, Vince D. Calhoun, Stefan Ehrlich, Lei Wang, Jessica A. Turner, and Nora I. Perrone Bizzozero. Meta gene set enrichment analyses link mir-137-regulated pathways with schizophrenia risk. Frontiers in Genetics, 6:147, 2015.

[23] Andrew D Yates, Premanand Achuthan, Wasiu Akanni, James Allen, Jamie Allen, Jorge Alvarez-Jarreta, M Ridwan Amode, Irina M Armean, Andrey G Azov, Ruth Bennett, Jyothish Bhai, Konstantinos Billis, Sanjay Boddu, José Carlos Marugán, Carla Cummins, Claire Davidson, Kamalkumar Dodiya, Reham Fatima, Astrid Gall, Carlos Garcia Giron, Laurent Gil, Tiago Grego, Leanne Haggerty, Erin Haskell, Thibaut Hourlier, Osagie G Izuogu, Sophie H Janacek, Thomas Juettemann, Mike Kay, Ilias Lavidas, Tuan Le, Diana Lemos, Jose Gonzalez Martinez, Thomas Maurel, Mark McDowall, Aoife McMahon, Shamika Mohanan, Benjamin Moore, Michael Nuhn, Denye N Oheh, Anne Parker, Andrew Parton, Mateus Patricio, Manoj Pandian Sakthivel, Ahamed Imran Abdul Salam, Bianca M Schmitt, Helen Schuilenburg, Dan Sheppard, Mira Sycheva, Marek Szuba, Kieron Taylor, Anja Thormann, Glen Threadgold, Alessandro Vullo, Brandon Walts, Andrea Winterbottom, Amonida Zadissa, Marc Chakiachvili, Bethany Flint, Adam Frankish, Sarah E Hunt, Garth IIsley, Myrto Kostadima, Nick Langridge, Jane E Loveland, Fergal J Martin, Joannella Morales, Jonathan M Mudge, Matthieu Muffato, Emily Perry, Magali Ruffier, Stephen J Trevanion, Fiona Cunningham, Kevin L Howe, Daniel R Zerbino, and Paul Flicek. Ensembl 2020. Nucleic Acids Research, 48(D1):D682–D688, 11 2019.

[24] Konstantinos Zagganas, Thanasis Vergoulis, Maria D. Paraskevopoulou, Ioannis S. Vlachos, Spiros Skiadopoulos, and Theodore Dalamagas. BUFET: boosting the unbiased mirna functional enrichment analysis using bitsets. BMC Bioinformatics, 18(1):399:1–399:8, 2017.

[25] Song Zhang, Jing Cao, Y. Megan Kong, and Richard H. Scheuermann. GO-Bayes: Gene Ontology-based overrepresentation analysis using a Bayesian approach. Bioinformatics, 26(7):905–911, 02 2010.

[26] Xinping Zhu, Masahisa Kudo, Xiangjie Huang, Hehuan Sui, Haishan Tian, Carlo M. Croce, and Ri Cui. Frontiers of microrna signature in non-small cell lung cancer. Frontiers in Cell and Developmental Biology, 9:771, 2021.

